# Closed-Loop Auditory Stimulation Reveals Differential Sleep Oscillatory Contributions to Memory in Healthy Older Adults

**DOI:** 10.64898/2026.08.17.745219

**Authors:** Saied Sabaghypour, Lawrence Oprea, Arthur S. Powanwe, Christine N. Moreau, Natasha Alfeche, Adrian M. Owen, Stefan Kohler, Lyle E. Muller, Laura J. Batterink

## Abstract

Sleep oscillations during non-rapid eye movement (NREM) sleep support memory consolidation but decline with age. Phase-locked auditory stimulation (PLAS) enhances slow-wave activity, yet its effects on distinct oscillatory components and memory in older adults remain unclear. Sixteen healthy older adults (≥60 years; mean age = 65.06 ± 3.53 years) participated in a randomized, single-blind, sham-controlled crossover study. Participants completed two stimulation nights and two sham nights in a sleep laboratory. Changes in slow oscillations (0.5–1.25 Hz), frontal theta (4–8 Hz), centrofrontal slow spindles (12–14 Hz), and centroparietal fast spindles (14–16 Hz) were compared between stimulation and sham conditions. Declarative memory was assessed using a word-pair recall task that required overnight retention, and broader cognitive performance was evaluated using the Creyos cognitive assessment battery. PLAS enhanced sleep oscillatory activity without altering sleep architecture. Compared with sham, stimulation increased slow oscillation, frontal theta, centrofrontal slow spindle, and centroparietal fast spindle power across both stimulation nights. Although word-pair recall did not improve at the group level, individual differences in stimulation-induced increases in fast spindle power were positively associated with individual differences in overnight memory improvement. Closed-loop auditory stimulation enhances multiple NREM oscillations in healthy older adults while preserving sleep architecture. Moreover, stimulation-induced increases in fast spindle activity track individual differences in overnight memory improvement, suggesting fast spindles as a physiological marker of successful sleep-dependent memory consolidation and a potential target for sleep-based neuromodulation in aging.

## Introduction

Sleep plays a critical role in brain health. Yet, sleep quality generally deteriorates with age (Corbo et al., 2023; Porter et al., 2015), and disrupted sleep leads to poorer memory and cognitive outcomes (Hyndych et al., 2025; Khan & Al-Jahdali, 2023). Slow wave sleep (SWS) in particular is known to be especially important for memory consolidation (Born, 2010; Brodt et al., 2023; Höller et al., 2024; Walker, 2009), as well as other cognitive processes such as executive function and attention (S. J. Schreiner et al., 2021; Stepan et al., 2021). According to the active systems consolidation model, during SWS, newly encoded memories are repeatedly reactivated and gradually reorganized in neocortical regions (Born & Wilhelm, 2012; Diekelmann & Born, 2010; Gilboa & Moscovitch, 2021; Goto, 2022). At the center of the active systems consolidation mechanism are slow oscillations (SOs), which occur at a frequency of ∼1 Hz and reflect alternating up-states (depolarization and elevated population firing) and down-states (hyperpolarization and relative neuronal silence) (Neske, 2016; Steriade et al., 2001). It is widely recognized that SO activity during SWS coordinates higher-frequency rhythms, with their up-phase coinciding with increased spindle activity, as reflected by increased power in the sigma band (12-16 Hz) (Marshall et al., 2003; Nir et al., 2011). Theta (4-8 Hz) power is similarly modulated by SO phase, with increases toward the down phase observed in scalp and intracranial EEG, supporting a role for theta-band events in the micro-architecture of NREM oscillatory sequences (Cox, van Driel, et al., 2014; Gonzalez et al., 2018a). From the perspective of active systems consolidation, these observations suggest that if SOs define windows of excitability and silence, then nesting of other frequency bands provides a timing mechanism that can implement hippocampo–neocortical dialogue (Klinzing et al., 2019; T. Schreiner et al., 2021). Accordingly, age-related reductions in SO activity, associated with prefrontal atrophy (Mander et al., 2013), may impair the coordination of sleep oscillations and contribute to reduced memory consolidation (Harand et al., 2012; Mander et al., 2017).

Spindles are a key component within this framework, which are increasingly recognized as heterogeneous rather than uniform. Spindles are divided into two subtypes based on their frequency and scalp distribution: centrofrontal slow spindles (12–14 Hz) and centroparietal fast spindles (14–16 Hz), each with distinct neural generators (Schabus et al., 2007a; Zeitlhofer et al., 1997a). At the systems level, slow spindles have been proposed to primarily reflect cortico-cortical interactions, whereas fast spindles involve thalamo-cortical communication (Doran, 2003), and are rooted in the precuneus and memory-relevant areas including the hippocampus (Anderer et al., 2001; Schabus et al., 2007b). In addition to their involvement in long-term potentiation, fast spindles temporally coordinate with hippocampal activity and are thought to facilitate information exchange between the hippocampus and the neocortex, underlying sleep-dependent memory consolidation (Buzsáki, 1996; Clemens et al., 2011). Relative to slow spindles, fast spindles are thus thought to play a more central role in memory consolidation (Saletin et al., 2011; van der Helm et al., 2011). Along with fast spindles, theta activity during NREM sleep may also contribute to memory consolidation (Jiang et al., 2017; T. Schreiner et al., 2018). Theta-band oscillations have been associated with the beneficial effects of targeted memory reactivation, in which learning-related cues (e.g., odors or sounds) are presented during NREM sleep to enhance memory consolidation (Lehmann et al., 2016; T. Schreiner et al., 2015, 2018). However, despite growing evidence, NREM theta is still not considered a central component in theoretical models. Understanding how spindles and theta activity support sleep-dependent plasticity is essential, as this may provide a foundation for approaches targeting sleep-related oscillations and memory consolidation.

Importantly, slow-wave and spindles decline with advancing age, and such reductions have been linked to impairments in sleep-dependent memory consolidation (Champetier et al., 2023; Fogel et al., 2012; Pace-Schott & Spencer, 2014). Phase-locked auditory stimulation (PLAS) is a promising, non-invasive approach that has been shown to enhance slow oscillatory and spindle activity during sleep in young adults (Nguyen et al., 2023; Xi et al., 2023; Zeller et al., 2023), and thus has become a focus of research aimed at improving sleep in older adults because of its potential to counteract age-related alterations in these sleep rhythms. PLAS algorithms deliver brief bursts of pink noise precisely time-locked to the up-phase of endogenous SOs (Krugliakova et al., 2026; Ngo et al., 2013; Ong et al., 2016a; Sabaghypour et al., 2026). In healthy young adults, PLAS enhances SO and associated spindle dynamic, and importantly, improve memory consolidation at the behavioral level (Leminen et al., 2017a; Ngo et al., 2013, 2015; Ong et al., 2016b; Wunderlin et al., 2021). In contrast, in older adults, PLAS has led to more mixed effects, with one study finding enhancements in SO and improved declarative memory performance (Papalambros et al., 2017), and others reporting null effects and even worsened declarative retention in some cohorts (Schneider et al., 2020; Wunderlin et al., 2023). Furthermore, most prior PLAS studies in older adults have primarily focused on measures of SOs (Lustenberger et al., 2022) and overall spindle power in relation to declarative memory performance (Papalambros et al., 2019), and derived their findings based on single-channel analysis (Papalambros et al., 2017, 2019). Given the heterogeneous nature of sleep oscillations, it is critical to characterize the effect of PLAS in older adults on all key sleep rhythms—namely SOs, slow spindles, fast spindles and theta—and to determine whether stimulation-related changes at the individual level influence memory consolidation as well as other cognitive domains beyond declarative memory.

Here, we investigated the effects of PLAS on NREM oscillatory activity using a counterbalanced crossover design in which each participant completed two consecutive stimulation nights and two consecutive sham nights. Specifically, we examined whether stimulation enhances SOs, theta, slow spindles and fast spindles across the scalp. We further explored whether individual differences in these oscillatory enhancements would predict overnight declarative memory consolidation. In addition, we investigated whether stimulation is related to subsequent performance across broader cognitive domains, including short-term memory, reasoning, concentration, planning, and global cognitive performance.

## Methods

### Participants

All prospective participants were initially screened using the Brain Health Assessment (BHA; Troyer et al., 2014), an online, self-administered cognitive assessment tool that has been shown to provide similar detection of aMCI (amnestic mild cognitive impairment) as a clinician-administered screener (MoCA; Paterson et al., 2021). Sixteen cognitively healthy older adults (aged ≥60 years; female = 12; mean age= 65.06, SD=3.53), as determined by a BHA z-score of 0.60 or below were selected and included in the present study, indicative of normal aging and relatively low risk for aMCI (a score of 0.64 and above is classified as higher risk of aMCI; Paterson et al., 2021). Exclusion criteria included (1) diagnosis of major psychiatric or neurological disorders, (2) moderate to severe symptoms of depression or anxiety, (3) diagnosed sleep disorders, (4) serious medical illness, (5) history of stroke or transient ischemic attack, (6) current or past alcohol or substance abuse, (7) history of seizures, (8) regular use of psychoactive or hypnotic medications or any medication known to alter sleep, (9) significant hearing loss or use of hearing aids (see details below), and (10) untreated moderate or severe sleep apnea, defined as an apnea-hypopnea index (Olson et al., 2003) (AHI) ≥15 events/hour based on a home sleep apnea test. Hearing status was assessed using self-report and an online screening measure. Participants indicated via yes/no questions whether they had good hearing and whether they used hearing aids. In addition, participants completed the Shoebox online hearing screening test (Shoebox Ltd). Participants were considered to have adequate hearing if they obtained a “good” score in at least one ear. All participants provided written informed consent before participation. The study protocol was approved by Western’s Health Sciences Research Ethics Board (HSREB) and conducted in accordance with the Declaration of Helsinki.

### Experimental design

This study employed a randomized, single-blind, sham-controlled crossover design conducted in a controlled sleep laboratory environment. Each participant completed five overnight sessions: an initial acclimation night, followed by two consecutive nights of either stimulation or sham, and then two consecutive nights of the alternate condition (see Figure 1a). The two experimental blocks were separated by a one-week washout period. The order of stimulation and sham blocks was counterbalanced across participants.

**Figure 1.**
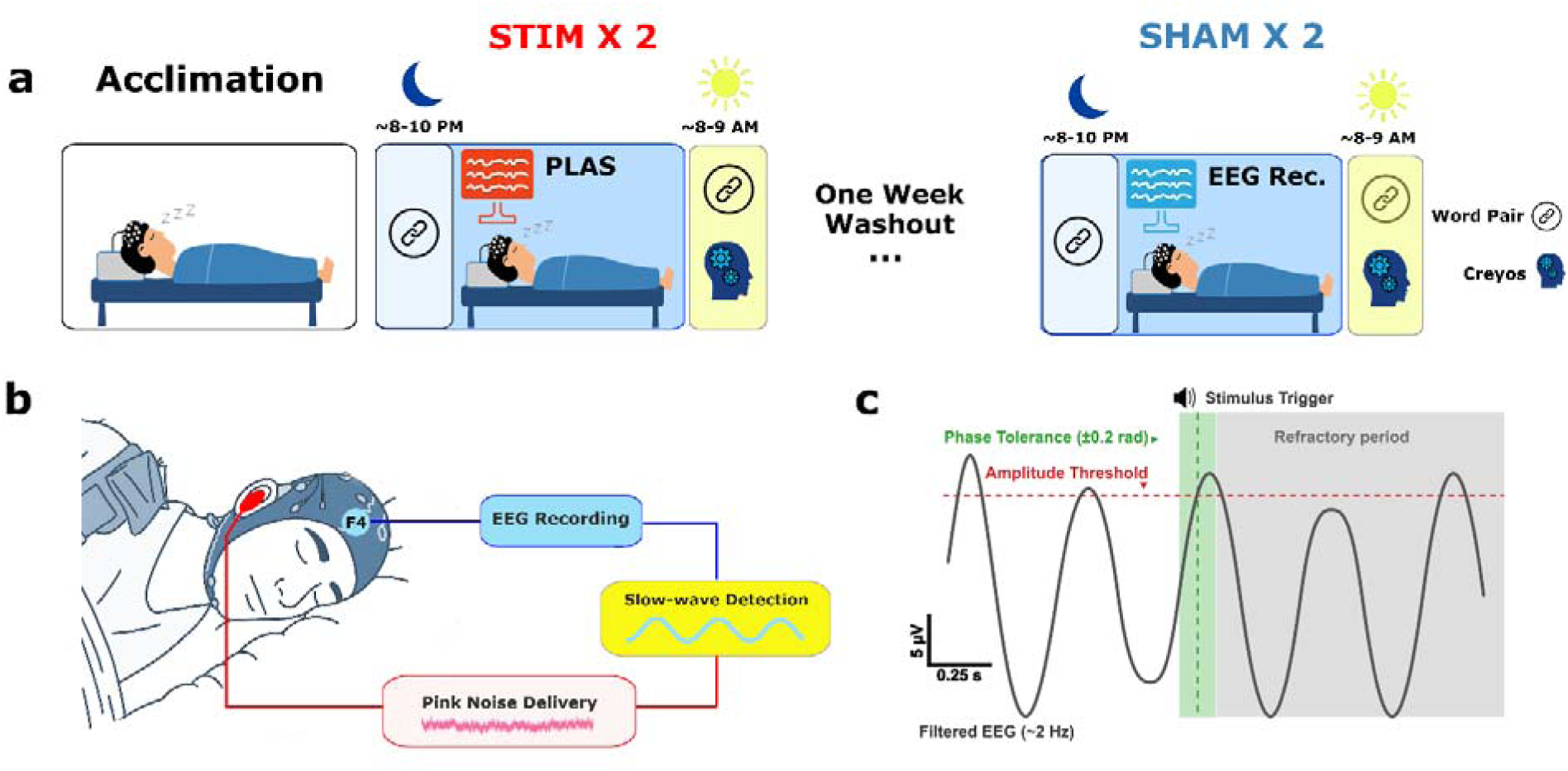
Overview of the experimental design and procedure. a) Schematic representation of the experimental procedure. Each night consisted of either STIM or SHAM (no stimulation) conditions, with block order (consisting of 2 consecutive nights) counterbalanced across participants. b) Illustration of the PLAS stimulation system. EEG signals were recorded in real time, SOs were detected online, and brief pink-noise stimuli (50 ms) were delivered at a targeted phase of the SO during NREM sleep. (c) Schematic illustration of the phase-targeting algorithm. Auditory stimulation was triggered only when the ongoing slow oscillation exceeded the amplitude threshold and its phase entered the predefined target window (±0.2 rad around the target phase). Following each stimulus, a refractory period (6s) prevented additional stimulation until the next eligible slow oscillation.

On each experimental night, participants arrived at the sleep laboratory approximately two hours before their habitual bedtime (∼8:00 PM). Participants first underwent setup for EEG recording, and then completed a word-pair learning task a followed by an immediate cued recall test to assess pre-sleep memory performance. Participants then received an overnight sleep opportunity while EEG was continuously recorded. During sleep, either PLAS (pink noise with 5 ms linear ramp-up/down; stimulation intensity individually titrated to each participant’s hearing threshold during pre-sleep setup and further adjusted online based on EEG indicators of responsiveness to avoid arousal, see Figure 1b) or “sham stimulation” was delivered using a real-time algorithm that targeted the positive peaks of detected SO during NREM sleep (see Figure 1c). In the sham condition, the algorithm ran as in the stimulation condition, but no sounds were delivered. Participants were blinded to conditions throughout the study. The following morning, participants completed a second delayed cued recall task for the previously learned word pairs, and the Creyos cognitive assessment battery (Hampshire et al., 2012)—a validated, web-based platform comprising multiple tasks that assess core cognitive domains including short-term memory, reasoning, attention, and executive function (Boa Sorte Silva et al., 2018; Gregory et al., 2016).

### EEG Recording

Sleep EEG data were collected using a 64-channel BrainVision system (Brain Products, Munich, Germany) with the online reference placed on the left mastoid (M1). Two electrodes were positioned near the outer corners of the eyes to record electrooculogram (EOG) activity, another pair was placed on the chin to monitor electromyographic (EMG) signals. Electrocardiography (ECG) signals were also recorded using standard electrode placement. All electrode impedances were maintained below 20 kΩ throughout setup. Continuous EEG signals were acquired at a sampling rate of 1000 Hz.

### Memory and cognitive tasks

Memory consolidation was assessed using a verbal paired-associate learning task (Figure 2a). Approximately 90 minutes prior to lights out, participants were shown 45 word pairs composed of moderately related nouns (e.g., garden–flower) (Ngo et al., 2013; Westerberg et al., 2012). Each word pair was displayed at the center of a computer screen for 4 seconds, with a 1-second interstimulus interval. Participants were instructed to memorize each pair. Immediately after the learning phase, participants completed an initial cued-recall test in which the first word of each pair was presented in random order. Participants were asked to recall the second word of the pair with a verbal response. Following each attempt, the correct pair was briefly displayed for one second, regardless of the participant’s response, providing feedback and an additional encoding opportunity. The next morning, participants completed a second cued-recall test, with words presented in a new random order. Memory consolidation was indexed as the change in number of correctly recalled words from the second (morning) session compared to the first (evening) session, calculated as ((Number of Correct Items at Morning Session − Number of Correct Items at Evening Session) / Number of Correct Items at Evening Session) × 100, with higher values indicating better overnight memory consolidation.

**Figure 2.**
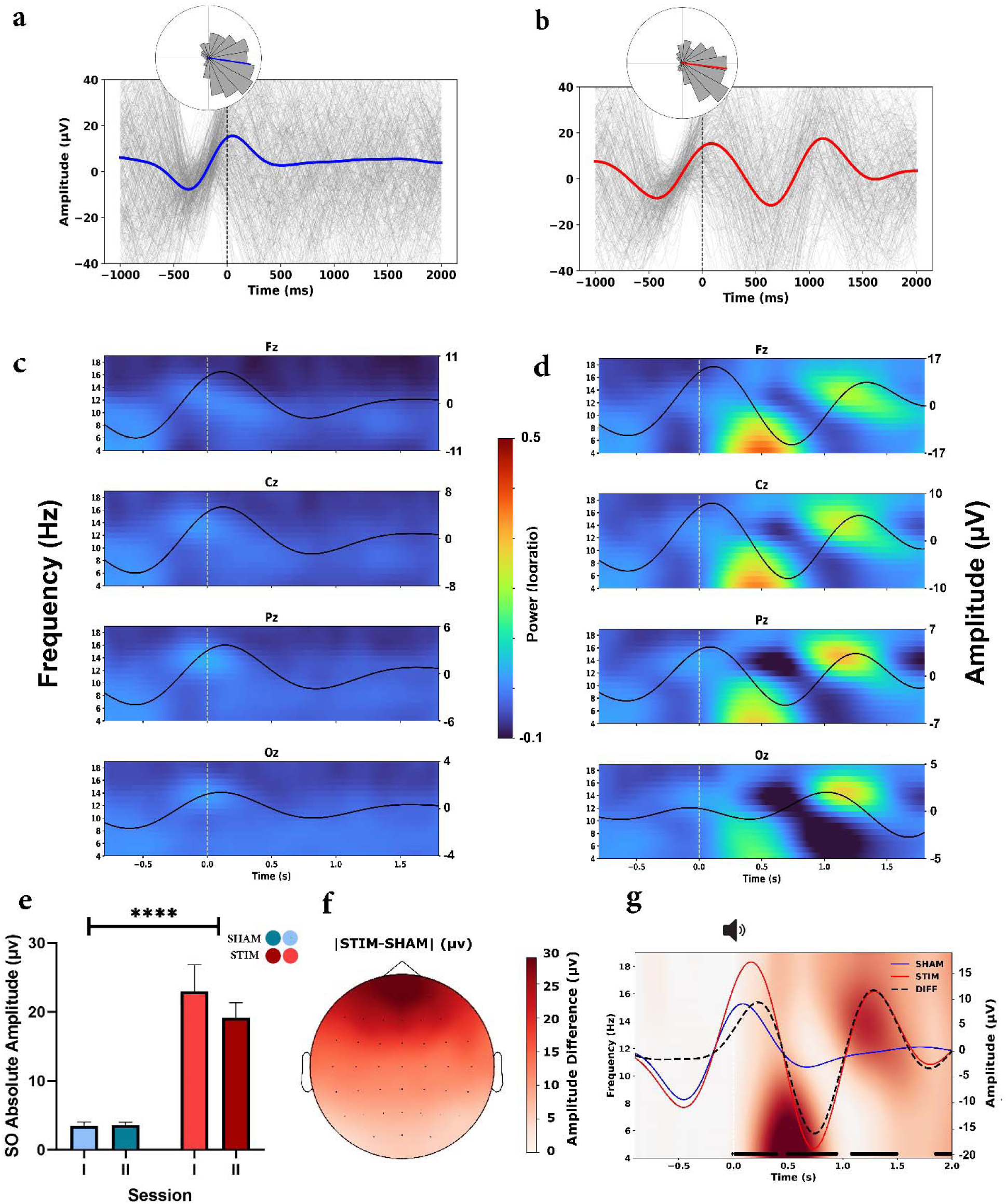
Channel-level, time–frequency and ERP representations for SHAM and STIM nights. (a–b) Sample ERPs at electrode F4 for SHAM (a) and STIM (b) conditions. Gray lines show individual trials for one representative session of STIM and SHAM, and colored traces indicate the mean response. Circular histograms below show phase at stimulus onset (blue: SHAM; red: STIM). (c–d) Time–frequency representations (TFRs) with overlaid ERPs for selected electrodes (Fz, Cz, Pz, Oz), shown for the mean of SHAM nights (c) and STIM nights (d). Black traces represent the ERP time course at each electrode. (e) Mean SO trough amplitude (absolute µV) for SHAM (blue) and STIM (red) conditions, computed within the SO band (0.5–1.25 Hz) (0–2 s window relative to stimulus onset). Error bars indicate SEM; asterisks denote significant condition effects. (f) Scalp topography of SO amplitude differences (STIM − SHAM), illustrating the spatial distribution of stimulation-related increases in SO amplitude. (g) Time– frequency representation with overlaid ERPs across frontal electrodes. Colored maps depict STIM − SHAM log-ratio power differences, with overlaid ERPs for STIM (red) and SHAM (blue). The black dashed trace represents the ERP difference waveform (STIM − SHAM), and black horizontal bars denote time intervals with significant ERP differences (p < .05, FDR-corrected). (TFR: time-frequency representation; ERP: event-related potential).

Cognitive performance was assessed using the Creyos cognitive assessment platform (Hampshire et al., 2012), which includes 12 brief, non-verbal tasks that tap a range of cognitive abilities (short term memory, reasoning, concentration and planning; see Supplementary Table S1 for the specific tasks contributing to each domain). This web-based testing system has been validated in large-scale studies and provides a reliable measure of global cognitive function (Wild et al., 2018). The outcome measures were calculated from participants’ performance across the 12 Creyos tasks. Consistent with previous methods (Boa Sorte Silva et al., 2021; Gill et al., 2015), task scores were z-transformed and then averaged to obtain domain-specific composite scores. These domain composites were also subsequently averaged to generate a global cognition score.

### Phased-locked stimulation

Real-time phase tracking of the incoming EEG data stream was achieved through a graphical software interface implemented in MATLAB 2024b (MathWorks, Natick, MA). An expert sleep technician monitored the EEG signal to determine NREM sleep onset, then activated real-time processing manually. EEG signal from electrode F4 referenced to M1 was streamed to the MATLAB interface for all phase-determinations steps. To mitigate edge artifacts, the raw data was symmetrically constant-padded then filtered, using a zero-phase 2nd order Butterworth bandpass filer (0.5 – 4 Hz). The instantaneous phase angle and amplitude envelopes were then extracted using the Hilbert transform.

The decision to deliver a phase-locked stimulus was continuously evaluated at each buffer update using a stringent set of temporal, amplitude, and phase criteria. To ensure stimulation was only delivered during periods of robust oscillatory activity, a representative amplitude was derived by calculating the median of the amplitude envelope over a recent, user-defined trailing temporal window (see Table 1 for parameters used). A stimulus trigger was dispatched only if three strict conditions were simultaneously met: (1) a minimum refractory period of six seconds had elapsed since the preceding stimulus to prevent temporal clustering; (2) the computed median amplitude exceeded a user-set microvolt threshold; and (3) the absolute circular distance between the current instantaneous phase and the target phase fell within a predefined accuracy tolerance. Upon satisfying these criteria, the pink noise was played. Audio processing and presentation were handled using Psychtoolbox-3 (Brainard, 1997), to ensure minimal latency in this system. The pink noise stimulus was delivered through an external sound card (Steinberg UR22C) into a mono channel studio monitor centered at the subject’s head. A trigger signal was sent to EEG amplifier to record the exact timing of the stimulus initiation. Finally, the precise timestamp, phase, and amplitude were appended to the MATLAB session logs for subsequent offline analysis.

**Table 1.**
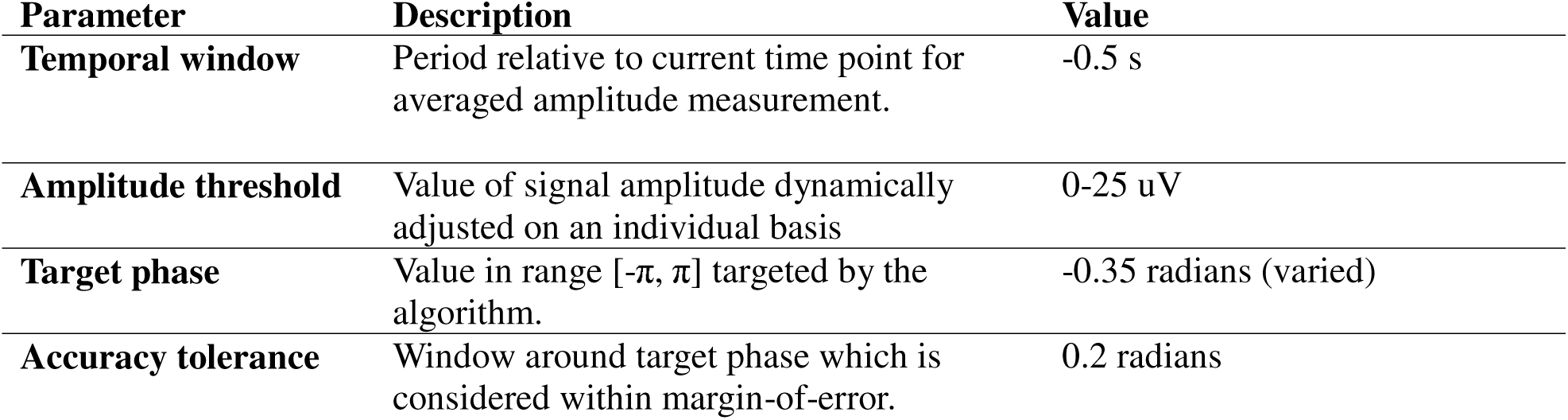
Parameters used in phase-detection algorithm.

Sleep stages and corresponding durations were classified for all participants using the YASA+MNE toolbox (v0.6.3), a machine-learning–based sleep staging algorithm trained and validated on approximately 30,000 hours of polysomnographic recordings (Vallat & Walker, 2021). In accordance with recommended preprocessing procedures (including central EEG, EOG, and EMG channels, downsampling to 100 Hz, and band-limited filtering between 0.4 and 30 Hz), the algorithm generated 30-s epoch hypnograms for each subject. From the resulting hypnograms, sleep architecture metrics were extracted, including total sleep time (TST), time spent in each sleep stage (N1, N2, N3, REM), and sleep efficiency (TST/time in bed × 100).

In addition, subjective sleep experience was assessed each morning using a brief questionnaire comprising multiple Likert-scale items (1–5). A subjective sleep score was calculated by averaging items reflecting overall sleep quality, restfulness, sleep depth, sleep continuity, and ease of sleep initiation and awakening (see Supplementary Materials for the list of questionnaire items).

### EEG preprocessing and event-related analysis

EEG analyses focused on NREM sleep for each night, which served as the basis for subsequent preprocessing and analyses. Data were band-pass filtered (FIR filter, 0.5-35 Hz, zero-phase non-causal FIR filter, firwin design with Hamming window). Cardiac artifacts were identified using independent component analysis (ICA) and removed based on their temporal correlation with the ECG signal. Noisy channels were identified through visual inspection and interpolated using spherical spline interpolation (Perrin et al., 1989). EEG data were then segmented into epochs time-locked to stimulation (or sham) onset (−1 to 2 s relative to stimulus onset). Epochs containing excessive artifacts were automatically removed using a voltage-range amplitude criterion across channels, defined as epochs whose voltage range exceeded the 95th percentile of all epoch voltage ranges (Mirjalili et al., 2021; Strunk et al., 2017). Visual inspection confirmed that rejected epochs predominantly reflected clear artifacts (e.g., movement-related noise). The remaining epochs were visually inspected, and any additional trials containing artifacts were also excluded from further analyses. In one participant, data from a single STIM night were excluded due to technical issues (excessive noise and missing stimulation triggers). Across all nights, an average of 421.4 ± 172.0 STIM/SHAM events were delivered per participant-night, of which 354.8 ± 146.6 epochs remained after preprocessing. On average, 15.8 ± 3.9% of epochs were rejected due to artifacts or noise.

Event-related potentials (ERPs) and time–frequency representations (TFRs) were computed from stimulation-locked epochs at each electrode and averaged within each participant and condition. To characterize SO dynamics, ERP signals were filtered in the SO range (0.5–1.25 Hz) using a zero-phase second-order Butterworth filter. After obtaining individual ERPs, group-level ERPs were then obtained by averaging across participants separately for SHAM and STIM conditions, and an ERP difference waveform (STIM − SHAM) was calculated.

To characterize spindle and theta activity, time–frequency decomposition was performed using Morlet wavelets (tfr_morlet) across frequencies from 4–20 Hz (wavelet cycles increasing linearly from 5 to 12 cycles) (Denis et al., 2021; Shinagawa et al., 2025). Power estimates were baseline corrected using a log-ratio normalization relative to the −1 to 0 s prestimulus interval. The time windows used to quantify theta and spindle activity were defined based on visual inspection of the grand-average TFR and prior literature (Cox, Korjoukov, et al., 2014; Leminen et al., 2017b), and included 300–600 ms for theta, and 800–1200 ms for spindle activity. Based on previous literature describing the spatial distribution of NREM theta and spindles, three regions of interest (ROIs) were defined to quantify stimulation-related oscillatory activity. Specifically, frontal electrodes were used to quantify theta (4-8 Hz) activity (Ferrara et al., 2002; Sprecher et al., 2016), and frontocentral and centroparietal electrodes were used to quantify slow (12-14 Hz) and fast spindle (14-16 Hz) activity, respectively (Cox et al., 2017a; Koupparis et al., 2013; Schabus et al., 2007c; see Supplementary Materials for the electrode composition of each ROI). Within each ROI, band-limited power was averaged across frequencies, time points, and channels to obtain power estimates per participant and condition. Confirmatory analyses further verified the spatial distribution of stimulation effects across the ROIs; detailed statistical outcomes are provided in the Supplementary Materials.

### Statistical Analysis

All statistical analyses were conducted in Graphpad Prism (GraphPad Software, Boston, Massachusetts USA, www.graphpad.com) and Python using custom scripts. Behavioral, EEG and sleep architecture variables were analyzed linear mixed-effects models (LME) and paired-samples t-tests as appropriate. Statistical significance was set at p < .05 (two-tailed) unless otherwise specified. Post-hoc tests were conducted to further explore significant main effects and interactions.

## Results

### Stimulation-related ERP replicate previously reported SO increases

As shown in the channel-level representations (Figure 2c–d), stimulation elicited a consistent increase in sleep oscillatory activity. An LME model including frontal channels (AF3, AFz, AF4, F4, F1, Fz, F2, F3, Fp1, Fp2) revealed a significant main effect of Condition, with larger (i.e., more negative) SO trough amplitudes during STIM compared to SHAM (β = 22.67, p < 0.001). There was no main effect of Session (p = 0.958) and no Condition × Session interaction (p = 0.304). To visualize condition differences in ERPs, paired-samples t-tests were performed at each time point on the subject-level filtered ERP signals, comparing STIM versus SHAM. Resulting p-values were corrected for multiple comparisons across time using the Benjamini–Hochberg FDR procedure (p< .05), and statistically significant intervals are visualized as black horizontal bars (Figure 2g).

### Across all three EEG measures, stimulation was associated with increases in spectral power relative to sham, with similar effects on both stimulation nights

LMEs were used to examine the effects of Condition (SHAM vs. STIM) and Session (Night 1 vs. Night 2 within each condition) on power in three EEG ROIs during NREM sleep (Figure 3a, b). All categorical predictors were sum-coded, such that fixed-effect estimates reflect deviations from the grand mean across conditions. A significant main effect of Condition indicated greater power during STIM compared with SHAM across nights for all three measures, including frontal theta (β = 0.157, SE = 0.010, z = 15.52, p < .001), centrofrontal slow spindle (β = 0.090, SE = 0.007, z = 13.18, p < .001), and centroparietal fast spindle (β = 0.136, SE = 0.008, z = 17.23, p < .001). There were no main effects of Session (all ps ≥ .056) and no Condition × Session interactions (all ps ≥ .847), indicating that stimulation effects were consistent across nights. Post hoc contrasts confirmed significantly greater power during STIM compared with SHAM on both Night 1 and Night 2 (all p < .001), with no significant differences between nights within either condition.

**Figure 3.**
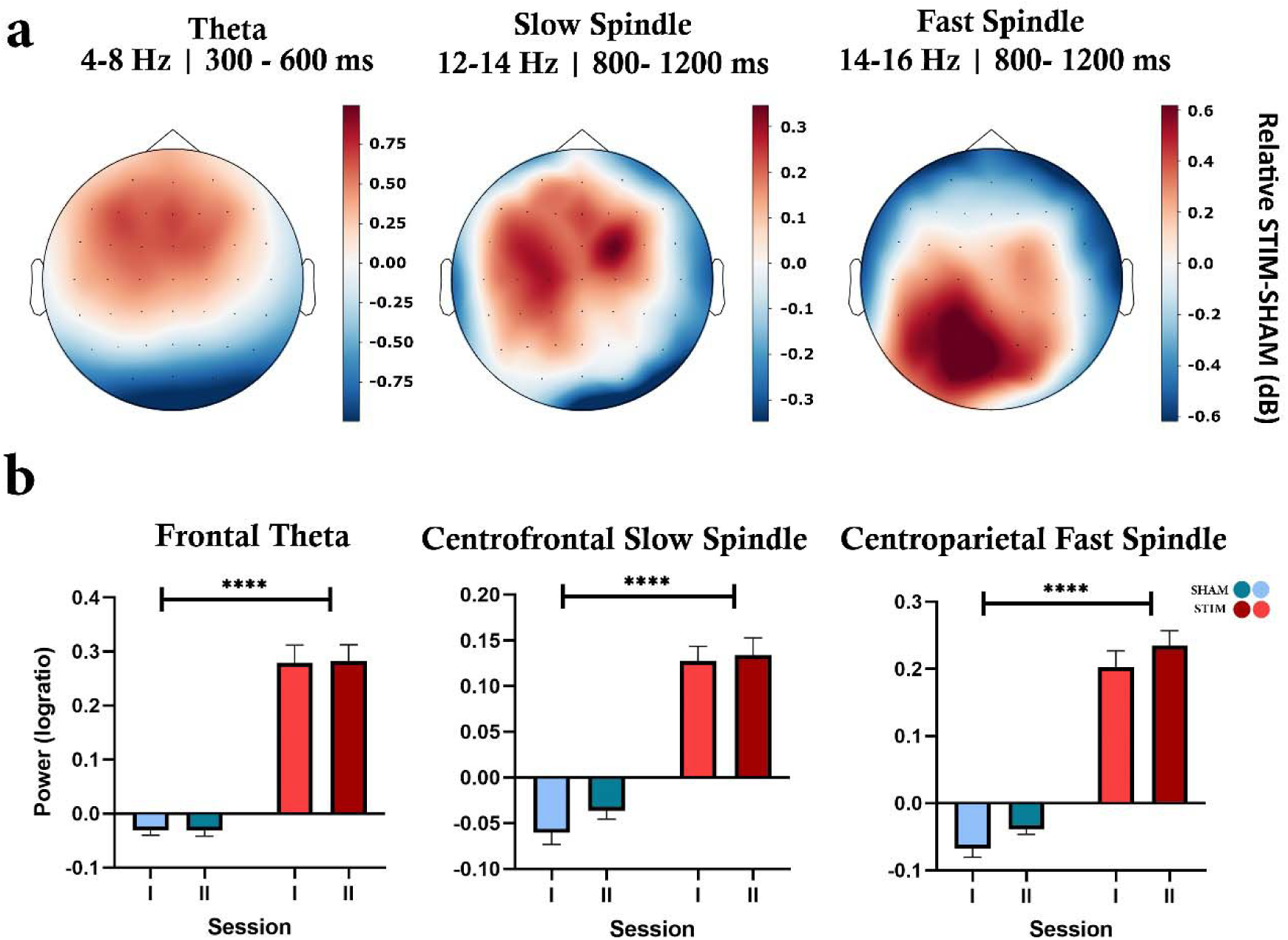
Stimulation-related modulation of NREM oscillatory activity. (a) Scalp topographies showing relative power differences (STIM – SHAM – mean centered) within predefined post-stimulus time windows for theta, slow and fast spindle. (b) Bar plots showing session-wise mean power (log ratio) for frontal theta, centrofrontal slow spindle, and centroparietal fast spindle during SHAM (blue) and STIM (red) conditions. Error bars indicate SEM; asterisks denote significant main effects of Condition.

### Mean sleep architecture measures did not differ between SHAM and STIM conditions

For pulses across trials, the mean instantaneous phase of pulse delivery was 341.1° (SD = 75.3°) for the STIM condition. Phase values computed in the same manner for SHAM trials yielded a mean phase of 350.1° (SD = 73.4°) Given that stimulation was targeted to the up-phase of the slow oscillation (approximately 320–340°), this indicates that pulse delivery was, on average, well aligned with the intended phase (Figure 4a, b).

**Figure 4.**
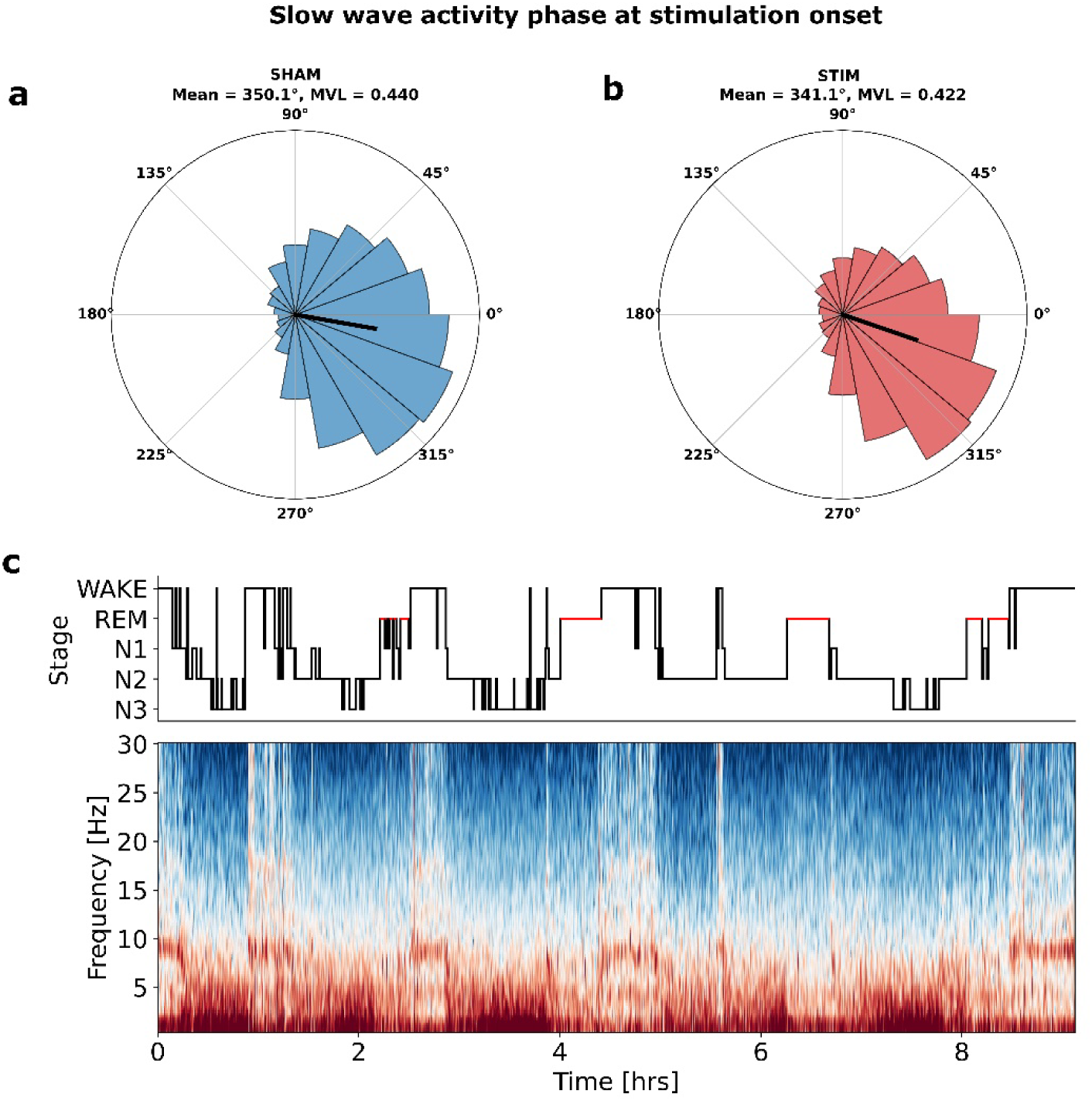
Slow-wave phase targeting. (a–b) Circular histograms showing the distribution of slow-wave activity (SWA) phase at stimulation onset for the SHAM (a) and STIM (b) conditions. Black arrows indicate the circular mean phase. (c) Representative hypnogram (top) and corresponding time–frequency spectrogram (bottom).

**Table 2.**
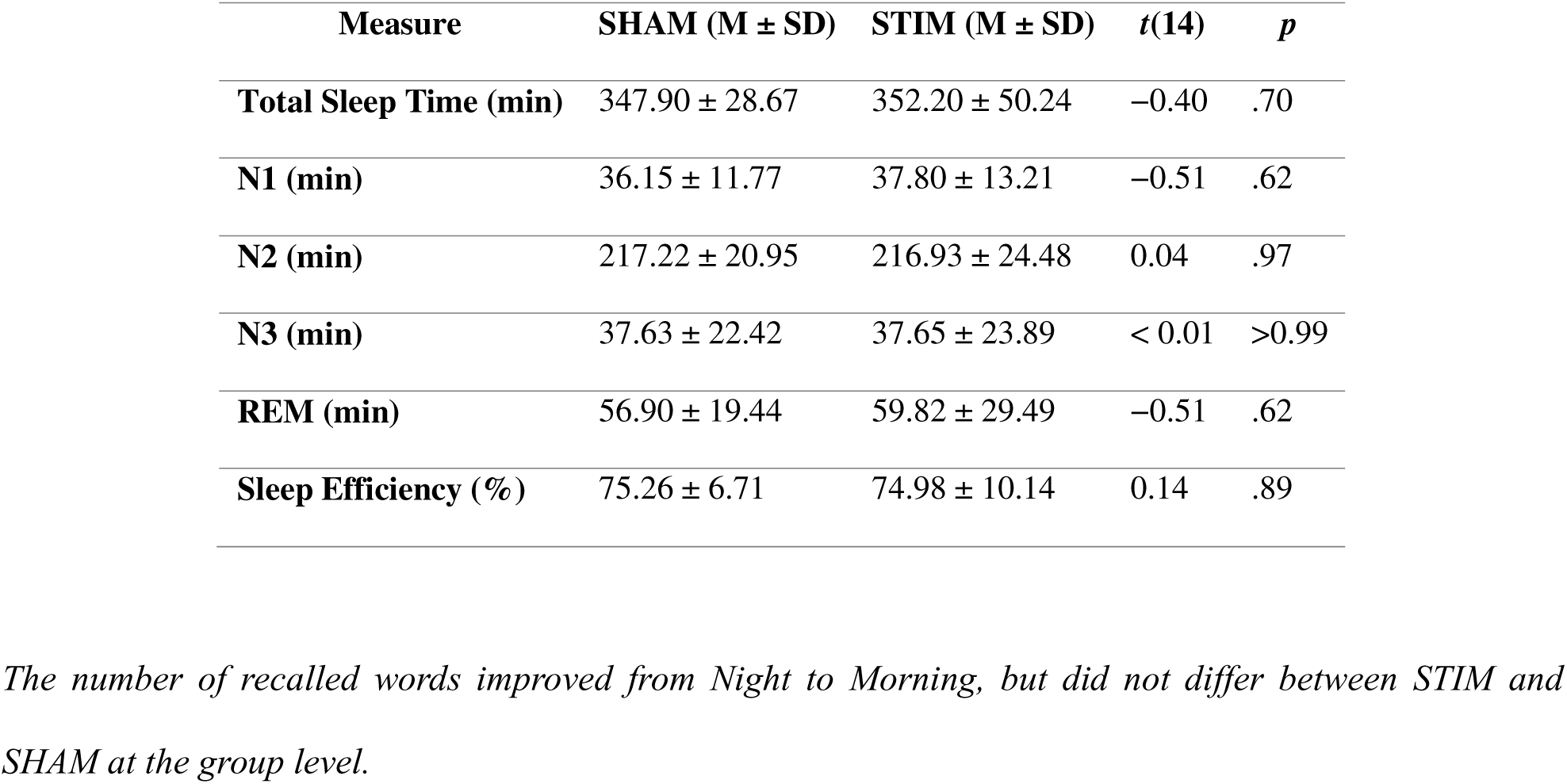
Mean Sleep Architecture Measures for SHAM and STIM Nights.

Mean sleep architecture measures did not differ between SHAM and STIM conditions. Paired-samples t-tests revealed no significant differences in total sleep time, time spent in any sleep stage (N1, N2, N3, REM), or sleep efficiency (all p values ≥ .62; Table 1). In addition, there was no significant difference in sleep scores between conditions, with participants reporting comparable levels of subjective sleep quality in the SHAM condition (M = 3.75) and the STIM condition (M = 3.62; t(14) = 0.94, p = .364, Cohen’s d = − 0.24). Together, these results indicate that stimulation did not alter overall sleep architecture.

### The number of recalled words improved from Night to Morning, but did not differ between STIM and SHAM at the group level

To examine changes in recall performance, we fitted an LME model using the absolute number of correctly recalled word pairs as the dependent variable. The model included Time (Night vs. Morning), Condition (SHAM vs. STIM), and Session (I vs. II) as fixed effects, with participant included as a random intercept. All categorical predictors were sum-coded, such that fixed-effect estimates reflect deviations from the grand mean across conditions. There was a significant main effect of Time, indicating higher recall in the Morning than at Night (β = −1.48, SE = 0.36, z = −4.14, p < .001). The main effects of Session (β = −0.14, p = .695) and Condition (β = −0.39, p = .276) were not significant, indicating no overall differences across sessions or between STIM and SHAM when averaged across other factors.

Memory improvement from Night to Morning was stronger in the first session across both STIM and SHAM conditions, compared to the second session (Time × Session (β = −0.70, *p* = .050). Follow-up comparisons examining overnight memory change (Night to Morning) within each experimental night revealed significant improvements in Session I for both SHAM (Δ = 3.81, SE = 1.44, z = 2.66, p = .032) and STIM (Δ = 4.94, SE = 1.44, z = 3.44, p = .002). No significant overnight change was observed in Session II for either SHAM or STIM (both p values ≥ .297; Figure 5b). Although not statistically significant, the overnight increase in recall was numerically larger for STIM, particularly for the first stimulation session. Furthermore, descriptive analyses indicated variability in individual response profiles. Numerically, 10 out of 16 participants exhibited greater overnight recall improvement in the STIM relative to the SHAM condition (see Supplementary Figure S1).

**Figure 5.**
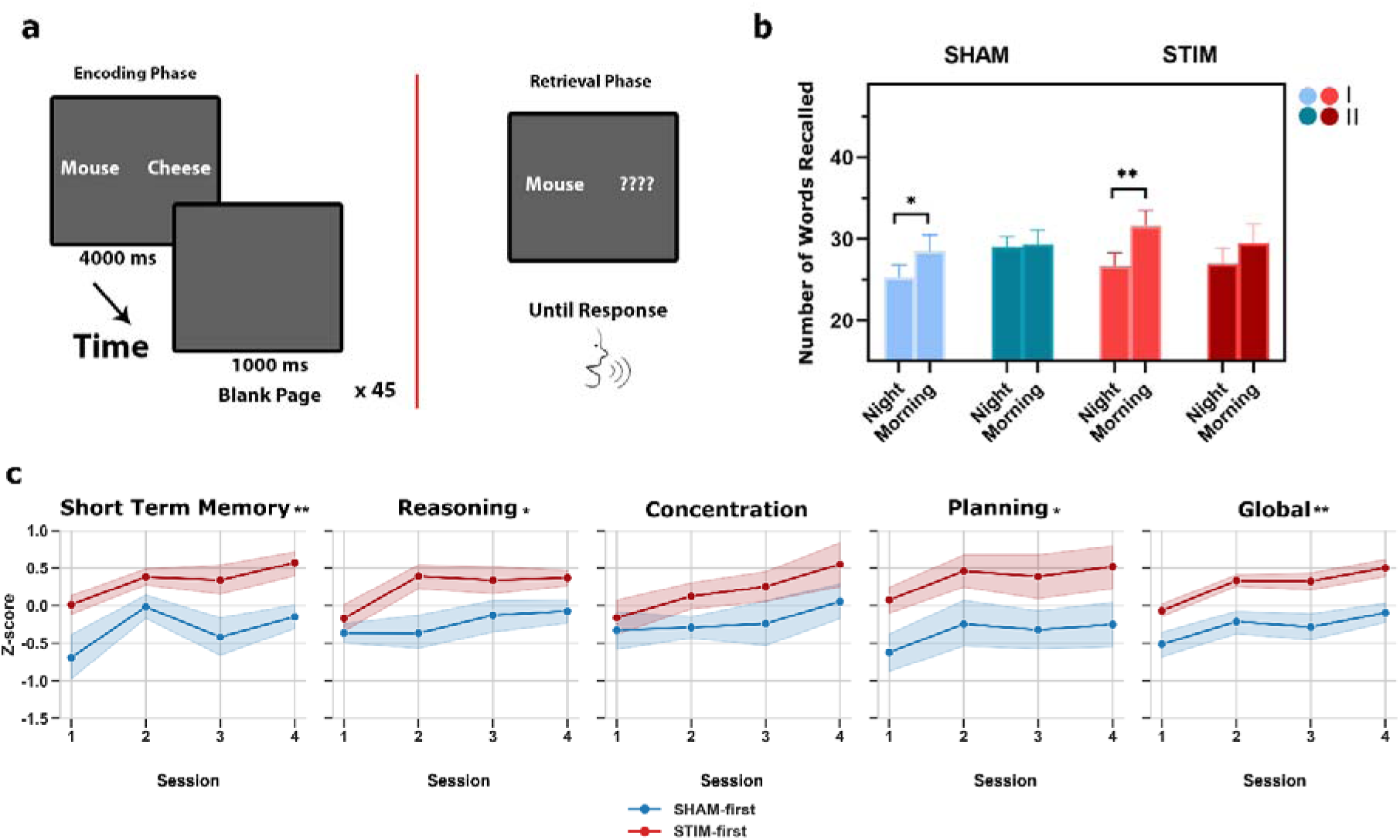
Behavioral performance across word pair and cognitive domains. (a) Schematic of the word-pair task. During encoding, word pairs were presented for 4s followed by a 1s blank screen (45 trials). During retrieval, cue words were presented until verbal response. (b) Mean number of recalled words at Night and Morning for SHAM and STIM sessions (I and II). (c) Composite cognitive performance across sessions as a function of counterbalancing order (SHAM-first vs. STIM-first). Participants who received STIM first showed overall higher performance in memory, reasoning, planning, and global cognition.

**Figure 6.**
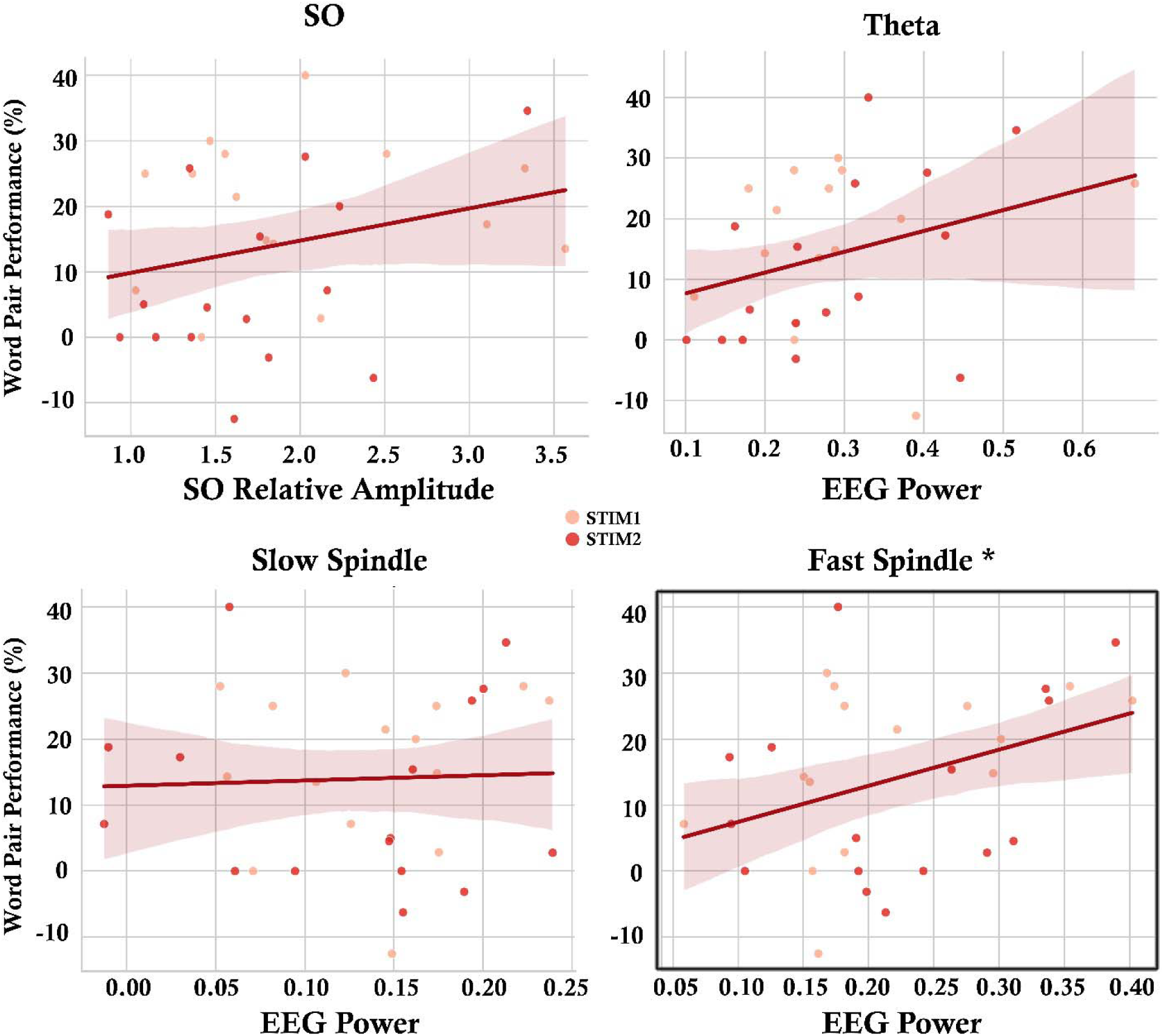
Relationships between stimulation-related EEG measures and overnight word-pair memory improvement during STIM nights. (a) SO relative amplitude, (b) frontal theta power, (c) centrofrontal slow spindle power, and (d) centroparietal fast spindle power. A significant association was found between fast spindle power and overnight memory improvement. Each point represents a participant-night (light and dark red indicate STIM1 and STIM2, respectively).

No other significant interactions were observed, including the Time × Condition (β = 0.39, p = .276), Condition × Session (β = −0.61, p = .090), or the three-way Time × Condition × Session interaction (β = −0.11, p = .761).

As an additional exploratory analysis, we averaged the overnight consolidation index (percentage change in recall from Night to Morning) across the two SHAM nights and across the two STIM nights for each participant to obtain a more stable estimate of condition-specific memory consolidation. Although mean percentage change was numerically greater during STIM (13.72%) than SHAM (8.66%), the difference was not statistically significant [paired t-test: t(15) = 1.22, p = .240] (see Figure S1).

### Stimulation did not alter overall performance across individual Creyos tasks

Performance on each Creyos task was analyzed using linear mixed-effects models with fixed effects of Condition (SHAM vs. STIM), Session (first vs. second night within each condition), and their interaction, and a random intercept for Participant (Supplementary Figure S2). Categorical predictors were sum-coded, such that fixed-effect estimates reflected deviations from the grand mean across conditions. To account for multiple testing across the 12 task-level analyses, p values were corrected using the Benjamini–Hochberg false discovery rate (FDR) procedure separately for each model effect (Condition, Session, and Condition × Session). After FDR correction, no task showed a significant main effect of Condition or a significant Condition × Session interaction (all ps ≥ .260).

In contrast, significant main effects of Session were observed for Paired Associates (p = .011), Spatial Planning (p = .011), Digit Span (p = .001), Polygons (p = .036), and Double Trouble (p = .033), reflecting higher performance in the second session than in the first, irrespective of stimulation condition. Full task-specific details are presented in Supplementary Table S1.

### Cognitive performance in memory, reasoning, planning, and global cognition was influenced by counterbalancing order, with higher scores observed in participants who received STIM first

As described above, cognitive performance was summarized into four domain composite scores (memory, reasoning, concentration, and planning), along with a global cognition composite. Given the repeated crossover design, we also evaluated whether overall cognitive performance differed according to the counterbalanced order of experimental conditions by conducting separate LMEs for memory, reasoning, concentration, planning, and global cognition Z-scores. All categorical predictors were sum-coded, such that fixed-effect estimates reflect deviations from the grand mean across conditions. The models included fixed effects of Condition (STIM vs. SHAM), Session (I vs. II), their interaction, and CounterbalanceOrder (STIM-first vs. SHAM-first), with a random intercept for Participant. Significant effects of CounterbalanceOrder emerged within four of these domains, including Memory (β = -0.32, p = .007), Reasoning (β = -0.23, p = .019), Planning (β = -0.36, p = .027), and Global Cognition (β = -0.27, p = .002), with higher overall performance observed in participants who received STIM first (Figure 5c; see Supplementary Materials for details). Overall session effects were additionally observed for Memory (β = - 0.19, p < .001) and Global Cognition (β = -0.13, p = .008), indicating improved performance from the first to the second night within each condition, with no evidence that this effect differed between STIM and SHAM. Concentration showed no significant effects. There were no significant main effects of Condition and no Condition × Session interactions.

### Stimulation-related fast spindle activity was positively associated with overnight memory improvement, with theta and SO amplitude showing trend-level associations

Pearson correlations were conducted to examine the relationship between stimulation-related EEG measures and overnight word-pair memory improvement within STIM nights. Frontal theta power showed a positive but non-significant association with memory improvement, r(29) = .314, p = .085, whereas no association was observed for centrofrontal slow spindle power, r(29) = .041, p = .827. In contrast, centroparietal fast spindle power was significantly positively associated with overnight memory improvement, such that higher fast spindle power during stimulation nights was related to greater word-pair memory gains, r(29) = .374, p = .038. SO relative amplitude showed no relation to overnight memory improvement, r(29) = .269, p = .143, though numerically greater SO relative amplitude was positively associated with memory gains.

## Discussion

Here, we investigated whether PLAS induces specific modulation of NREM oscillatory activity in healthy older adults and whether such neural effects lead to improvements in memory consolidation and broader cognitive performance. Using a randomized crossover design with two stimulation and two sham nights per participant, we aimed to extend prior work by differentiating slow and fast spindle subtypes and by incorporating the potential role of NREM theta oscillations.

Our closed-loop algorithm achieved accurate phase targeting, delivering auditory pulses approximately 10–20° before the peak of the endogenous slow-wave upstate. This temporal precision is particularly encouraging given the reduced SWA typically observed in older adults (Edwards et al., 2010; Lafrenière et al., 2023), which can make accurate phase targeting more challenging (Navarrete et al., 2020; Schneider et al., 2020). Furthermore, our PLAS algorithm elicited modulation of NREM oscillatory dynamics. In terms of SO (0.5-1.25 Hz) activity, we replicated prior reports showing that PLAS enhances SO-related responses when delivered at the up-phase of endogenous slow waves (Navarrete et al., 2020; Papalambros et al., 2017). Beyond the SO range and across both stimulation nights, spectral power in the theta and spindle bands was reliably enhanced relative to sham. Theta modulation emerged early (300 – 600 ms) following stimulation, whereas spindle responses occurred later (800 – 1200 ms), indicating a temporally ordered oscillatory response (Gonzalez et al., 2018a; Leach et al., 2024). This SO-linked increase in theta and spindle has been consistently reported, both after acoustic stimulation and during spontaneous slow waves (Cox, van Driel, et al., 2014; Gonzalez et al., 2018b; Klinzing et al., 2016; Mölle et al., 2011). Interestingly, similar time–frequency dynamics were observed in a TMS study, where single cortical pulses elicited increases in theta and spindle power following the evoked SO response (Bergmann et al., 2012). Beyond this temporal pattern, stimulation effects also showed frequency-specific topographies: theta increases were predominantly frontal, slow spindle effects were frontocentral, and fast spindle enhancements were centroparietal, consistent with known spatial distinctions between spindle subtypes (Cox et al., 2017b; Schönwald et al., 2012; Werth et al., 1997; Zeitlhofer et al., 1997b).

At the group level, we did not observe an overall significant difference in word pair memory performance between conditions in our sample (Schneider et al., 2020; Wunderlin et al., 2023). Yet, a majority of participants (10 out of 16) showed greater overnight improvement under STIM relative to SHAM (Supplementary Figure S1). This pattern may reflect individual differences in the integrity and functional organization of neural systems that support sleep-dependent memory consolidation. In older adults, variation in medial temporal lobe network and in sleep-related consolidation physiology has been linked to interindividual differences in overnight memory retention (Chappel-Farley et al., 2025). Such variability may, in turn, reflect differences in the capacity of these systems to respond to external modulation. In this respect, consolidation may be preferentially enhanced in individuals whose relevant neural networks retain sufficient plasticity (Boroshok et al., 2022; Stamps & Krishnan, 2014).

Consistent with this idea, individuals who exhibited greater stimulation-related increases in centroparietal fast spindle activity showed the largest gains in overnight word-pair memory; in other words, greater neural sensitivity to PLAS was linked to memory consolidation at the behavioural level. In contrast, slow spindle activity was unrelated to memory performance, whereas frontal theta power and SO relative amplitude showed positive but non-significant associations with memory improvement. These findings point to the importance of considering the potentially complementary contributions of stimulation-induced oscillatory dynamics to sleep-dependent memory consolidation. The selective association between fast spindle activity and memory performance converges with prior evidence suggesting that fast spindles play a particularly important role in systems consolidation during sleep (Barakat et al., 2011; Denis et al., 2021; Groch et al., 2017; Hahn et al., 2019; McDevitt et al., 2017; Mölle et al., 2011; Tamminen et al., 2010). Fast spindle activity has specifically been linked to increased hippocampal hemodynamic activity and the occurrence of hippocampal sharp-wave ripples (Clemens et al., 2011; Schabus et al., 2007c), and this relationship may represent a mechanism facilitating the reorganization of hippocampal–neocortical memory representations (Marshall et al., 2003; Wierzynski et al., 2009). Within this framework, greater stimulation-related fast spindle enhancement may reflect more effective coordination of hippocampal–neocortical communication during sleep. Importantly, these individual differences were not explained by baseline sleep oscillatory activity. Supplementary analyses showed that baseline SO, theta, and fast sigma power did not predict stimulation-related memory benefit (see Supplementary Materials), and baseline SO and fast sigma power were also unrelated to the magnitude of stimulation-induced neural enhancement. Interestingly, lower baseline theta power was associated with larger stimulation-induced theta enhancement (Supplementary Figure S6), suggesting that individuals with relatively reduced endogenous theta activity may retain greater capacity for PLAS-induced theta modulation. This finding should, however, be interpreted cautiously given the modest sample size and requires replication.

On the Creyos task, participants who received STIM first showed higher performance across memory, reasoning, planning, and global cognition (Figure 5), and this advantage persisted across subsequent SHAM nights. These results suggest that when effective, stimulation may induce plastic changes that could outlast the initial intervention (Richmond et al., 2014; Talsma et al., 2017). Nevertheless, although this observation of enduring effects in several cognitive domains is of potential theoretical and practical importance, such interpretation warrants caution, as the overall sample was split based on stimulation versus sham order. Hence, although participants were randomly assigned to counterbalance order, with the current sample size there is still a possibility of idiosyncratic pre-existing group differences.

Although theta power and SO relative amplitude did not reach statistical significance, both showed positive trend-level relationships with overnight memory improvement. During wakefulness, theta activity emerges during tasks requiring sustained processing (Kahana et al., 1999; Raghavachari et al., 2006). If theta serves a comparable organizing function during NREM sleep, it may structure cortical information processing prior to the down-state and facilitate the selection of relevant hippocampal traces for subsequent reactivation (Gonzalez et al., 2018b), whereas SOs provide the large-scale temporal framework that groups spindle and ripple activity during NREM sleep (Clemens et al., 2007; Niethard et al., 2018; Oyanedel et al., 2020). Within this framework, we may speculate that earlier theta may contribute to triggering the fast spindle-ripple sequence that underlies systems consolidation. Because hippocampal replay during sharp-wave ripples arrives at the cortex during the down-to-up transition as spindle activity emerges (Gonzalez et al., 2018b; Jiang et al., 2017; Maingret et al., 2016), stimulation-induced theta enhancement may reflect a heightened readiness of cortical networks to engage in this coordinated memory processing cascade. Thus, rather than acting independently, theta and fast spindle activity may represent temporally structured and functionally interdependent components of a broader consolidation mechanism (see Supplementary Materials Figure S3). The absence of association between slow spindle activity and memory improvement might support the notion that slow and fast spindles may play dissociable functional roles during sleep (Saletin et al., 2011; van der Helm et al., 2011). Together, these findings suggest that stimulation-induced enhancement of fast spindle activity may represent one of the most behaviorally relevant electrophysiological markers of successful PLAS-related memory consolidation.

## Conclusions

In summary, our PLAS algorithm reliably modulated sleep oscillatory dynamics during NREM sleep in healthy older adults. Across two stimulation nights, PLAS enhanced theta and spindle power without altering overall sleep architecture. In addition, stimulation-related increases in centroparietal fast spindle activity scaled with overnight improvement in word-pair recall. The findings of the present study suggest several directions for future work. First, larger samples will be necessary to clarify the relationship between stimulation-induced oscillatory modulation and behavioral outcomes. The absence of a group-level memory effect in the current study, despite clear neural modulation and declarative memory improvement in most participants, suggests inter-individual variability in responsiveness to PLAS. Future work should therefore aim to identify factors that predict behavioral responsiveness to stimulation.

Because aging is accompanied by reductions in SWA and sleep-dependent memory consolidation, interventions designed to enhance sleep oscillations may represent a non-invasive strategy for maintaining cognitive health across the lifespan. Identifying responsive individuals and assessing long-term effects will be critical steps toward translating sleep-based neuromodulation approaches into clinical and real-world settings.

## Data Availability

Data reported in this article are available from the corresponding author upon request.

