## Supplementary Materials for "Closed-Loop Auditory Stimulation Reveals Differential Sleep Oscillatory Contributions to Memory in Healthy Older Adults"

**Word Pair Task.** To further evaluate the consistency of the behavioral findings across sessions, we conducted exploratory analyses of the overnight memory consolidation index, calculated as the percentage change. First, consolidation scores were averaged across the two sham nights and the two stimulation nights (results reported in main text). We then examined consolidation separately for Session I and Session II and visualized individual participant trajectories across conditions. Exploratory paired comparisons between conditions within each session revealed no significant differences in overnight consolidation during either Session I, t(15) = -1.40, p = .182, or Session II, t(15) = -0.59, p = .562.

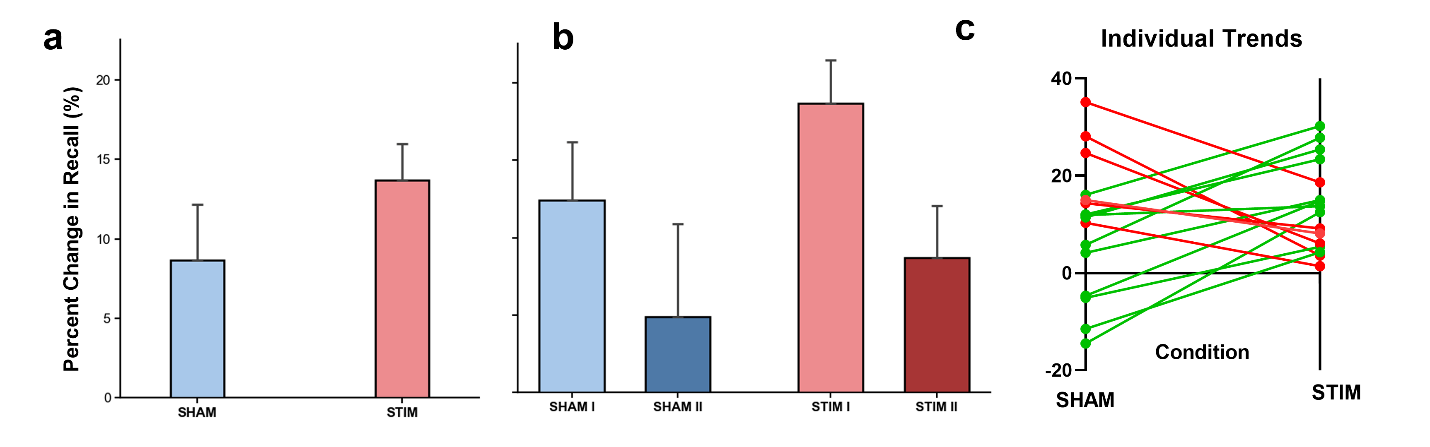

**Figure S1. Mean word-pair recall performance.** Exploratory analyses of overnight memory consolidation. (a) Mean overnight memory consolidation (percent change in word-pair recall from evening to morning) averaged across the two sham nights and the two stimulation nights for each participant. (b) Mean overnight memory consolidation shown separately for Session I and Session II under SHAM and STIM conditions. (c) Individual participant trajectories illustrating changes in overnight memory consolidation between SHAM and STIM after averaging across the two sessions; green lines indicate participants who showed greater consolidation during STIM than SHAM, whereas red lines indicate greater consolidation during SHAM than STIM..

**Creyos Tasks**

The twelve Creyos cognitive tasks and their corresponding cognitive domains are described in detail in Table S1.

Table S1. Description of the Cambridge Brain Sciences Cognitive Battery

| Task Name | Cognitive Domain | Brief Description |
| --- | --- | --- |
| 1. Monkey Ladder | Memory | Numbered squares appear simultaneously at random locations within an invisible 5×5 grid. After a delay, the numbers disappear and participants click the blank squares in ascending numerical order. Difficulty adapts trial-by-trial. |
| 2. Grammatical Reasoning | Reasoning | Participants determine whether a written statement correctly describes a pair of shapes displayed on the screen. Performance is time-limited and adaptive. |
| 3. Double Trouble | Reasoning | A colored word is displayed in incongruent or congruent ink. Participants identify which option matches the ink color of the word. Difficulty adapts with performance. |
| 4. Odd One Out | Reasoning | A 3×3 grid of patterned shapes is shown. Participants identify the item that violates an underlying rule relating color, shape, or number. |
| 5. Spatial Span | Memory | A sequence of squares in a 4×4 grid flashes sequentially. Participants reproduce the sequence in the same order. Sequence length adapts with performance. |
| 6. Rotations | Concentration | Two grids of colored squares are shown; one is rotated by multiples of 90°. Participants judge whether the grids are identical. |
| 7. Feature Match | Concentration | Two grids of abstract shapes are presented. Participants determine whether they are identical or differ by one element. |
| 8. Digit Span | Memory | A sequence of digits is presented sequentially. Participants reproduce the sequence using an on-screen keypad. Sequence length adapts across trials. |
| 9. Spatial Planning | Planning | Nine numbered beads are arranged on a tree-shaped frame. Participants reposition beads to achieve ascending numerical order using as few moves as possible. |
| 10. Paired Associates | Memory | Boxes open sequentially to reveal objects in random grid locations. Participants later recall the correct box location for each object. |
| 11. Polygons | Concentration | Overlapping polygons are displayed. Participants identify whether a target polygon matches one of the interlocking shapes. |
| 12. Spatial Search | Planning | Participants search for hidden tokens in boxes arranged on a grid, avoiding previously searched locations. Difficulty adapts across trials. |

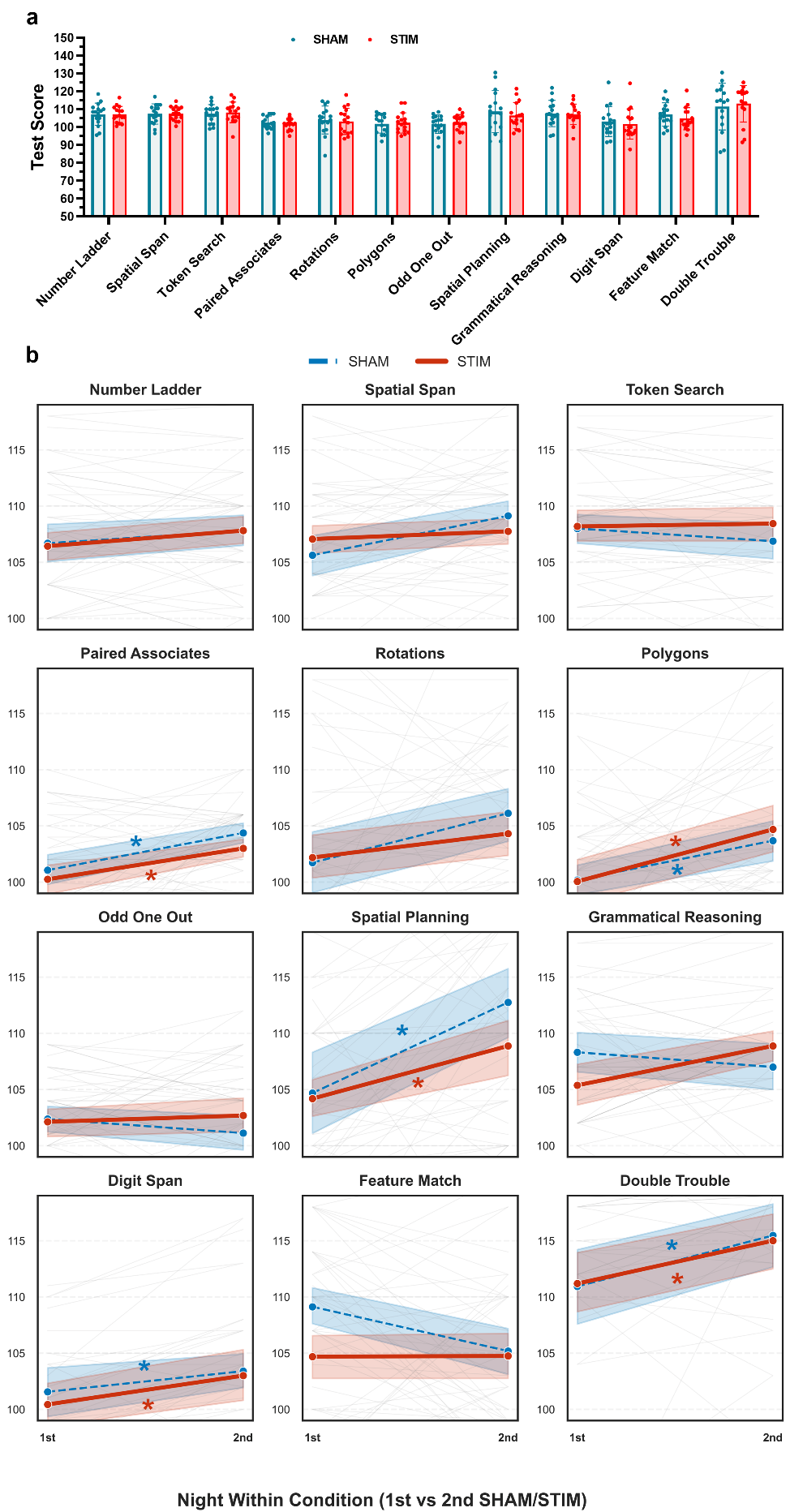

**Figure S2. Creyos cognitive performance across SHAM and STIM conditions.** (a) Mean task scores for SHAM and STIM. (b) Task-specific trajectories across the first and second administration (Session Within Condition), with individual participant lines in gray and condition means in color. A significant Condition × Session interaction emerged only for Grammatical Reasoning.

**LME with SessionWithinCondition and Counterbalance for Domains.** LMEs were conducted for memory, reasoning, concentration, planning, and global cognition Z-scores. All categorical predictors were sum-coded, such that fixed-effect estimates reflect deviations from the grand mean across conditions. The models included fixed effects of Condition (STIM vs. SHAM), Session (I vs. II), their interaction, and CounterbalanceOrder (STIM-first vs. SHAM-first), with a random intercept for Participant.

For Memory, there was no main effect of Condition, β = 0.05, SE = 0.04, z = 1.06, p = .289, and no Condition × SessionWithinCondition interaction, β = −0.03, SE = 0.04, z = −0.77, p = .443. However, a significant main effect of SessionWithinCondition was observed, β = −0.19, SE = 0.04, z = −4.37, p < .001, along with a significant effect of CounterbalanceOrder, β = −0.32, SE = 0.12, z = −2.70, p = .007.

For Reasoning, there was no main effect of Condition, β = −0.01, SE = 0.05, z = −0.12, p = .908, no main effect of SessionWithinCondition, β = −0.08, SE = 0.05, z = −1.54, p = .123, and no Condition × SessionWithinCondition interaction, β = 0.07, SE = 0.05, z = 1.40, p = .161. A significant effect of CounterbalanceOrder was present, β = −0.23, SE = 0.10, z = −2.35, p = .019.

For Concentration, there were no significant effects of Condition, β = 0.05, SE = 0.07, z = 0.76, p = .447, SessionWithinCondition, β = −0.11, SE = 0.07, z = −1.70, p = .089, CounterbalanceOrder, β = −0.20, SE = 0.13, z = −1.48, p = .138, or their interaction, β = 0.03, SE = 0.07, z = 0.46, p = .643.

For Planning, there was no main effect of Condition, β = 0.01, SE = 0.06, z = 0.15, p = .877, no effect of SessionWithinCondition, β = −0.12, SE = 0.06, z = −1.91, p = .056, and no Condition × SessionWithinCondition interaction, β = −0.01, SE = 0.06, z = −0.11, p = .912. A significant effect of CounterbalanceOrder was observed, β = −0.36, SE = 0.16, z = −2.21, p = .027.

For Global cognition, there was no main effect of Condition, β = 0.03, SE = 0.03, z = 0.84, p = .403, and no Condition × SessionWithinCondition interaction, β = 0.01, SE = 0.03, z = 0.43, p = .670. In contrast, significant main effects were found for SessionWithinCondition, β = −0.13, SE = 0.03, z = −4.19, p < .001, and CounterbalanceOrder, β = −0.28, SE = 0.09, z = −3.12, p = .002.

**Analysis of ROI-specific stimulation effects**

The channel composition of each region of interest (ROI) was defined as follows: frontal (theta; AF3, AFz, AF4, F3, F1, Fz, F2, F4), frontocentral (slow spindle; FC3, FC1, FCz, FC2, FC4, C3, C1, Cz, C2, C4), and centroparietal (fast spindle; CP3, CP1, CPz, CP2, CP4, P3, P1, Pz, P2, P4). To confirm the spatial distribution of stimulation-related oscillatory changes described in the main analyses, an LME model was fitted. The model examined the effects of Condition (STIM vs. SHAM), ROI (Frontal, Frontocentral, Centroparietal), and Frequency Band (theta, slow spindle, fast spindle) on EEG power (see also Figure 5a). The model revealed a main effect of Condition, such that stimulation was associated with higher power relative to sham (β = 0.215, SE = 0.017, z = 12.87, p < .0001). There was also a significant main effect of Band, with theta power exceeding fast-spindle power in the reference ROI (Frontal) (β = 0.042, SE = 0.017, z = 2.49, p = .013), whereas slow spindle did not differ from fast spindle (p = .39). No main effects of ROI were observed (all ps > .24). Interestingly, stimulation effects differed as a function of both ROI and frequency band. A significant Condition × ROI interaction indicated that the stimulation effect was stronger in centroparietal than frontal regions for fast spindle power (β = 0.061, SE = 0.024, z = 2.57, p = .0103). In addition, a significant Condition × Band interaction showed that stimulation enhanced theta power more strongly than fast spindle in frontal regions (β = 0.010, SE = 0.024, z = 4.22, p < .001). Importantly, these effects were further qualified by significant Condition × ROI × Band interactions, indicating that the spatial distribution of stimulation effects differed across frequency bands. Specifically, relative to fast spindle, stimulation-related increases in slow spindle were attenuated in centroparietal regions (β = −0.066, SE = 0.033, z = −1.99, p = .046), and stimulation-related theta increases were reduced in centroparietal compared to frontal regions (β = −0.123, SE = 0.033, z = −3.67, p < .001).

**Association between stimulation-related changes in frontal theta and spindle power.**

To examine whether stimulation-related changes in oscillatory activity were associated across frequency bands, participant-level difference scores (STIM–SHAM) were computed for theta, slow spindle, and fast spindle power. There was no significant association between stimulation-related changes in theta and slow spindle power, r(14) = .26, p = .660. In contrast, stimulation-related increases in theta were significantly positively correlated with increases in fast spindle power, r(14) = .59, p = .034.

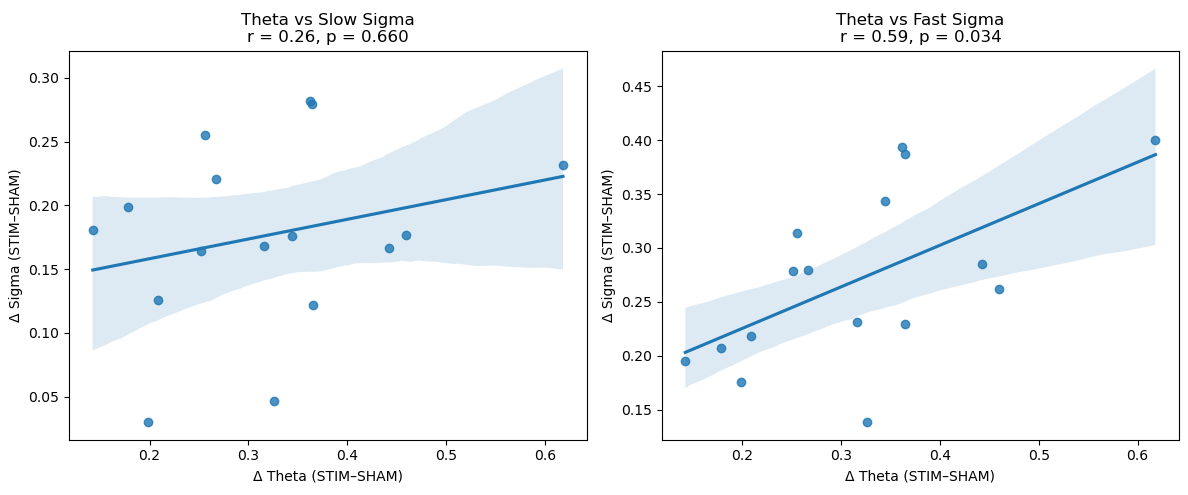

**Figure S3.** Correlation between stimulation-induced changes (STIM–SHAM) in theta and spindle power

**Subjective Sleep Questionnaire Items.**

Participants completed a brief morning questionnaire assessing subjective sleep experience. Responses were provided on a 5-point Likert scale (1–5). The items included in the subjective sleep score are listed below.

| Question | Response Scale |
| --- | --- |
| How did you sleep? | 1 = very poorly, 5 = very well |
| Did you feel refreshed after you arose this morning? | 1 = not at all, 5 = completely |
| Did you sleep soundly? | 1 = very restless, 5 = very soundly |
| Did you sleep throughout the time allotted for sleep? | 1 = woke up much too early, 5 = slept through the night |
| How easy was it for you to wake up? | 1 = very easy, 5 = very difficult |
| How easy was it for you to fall asleep? | 1 = very easy, 5 = very difficult |

**Channel-wise time–frequency representations of activity in SHAM and STIM conditions**

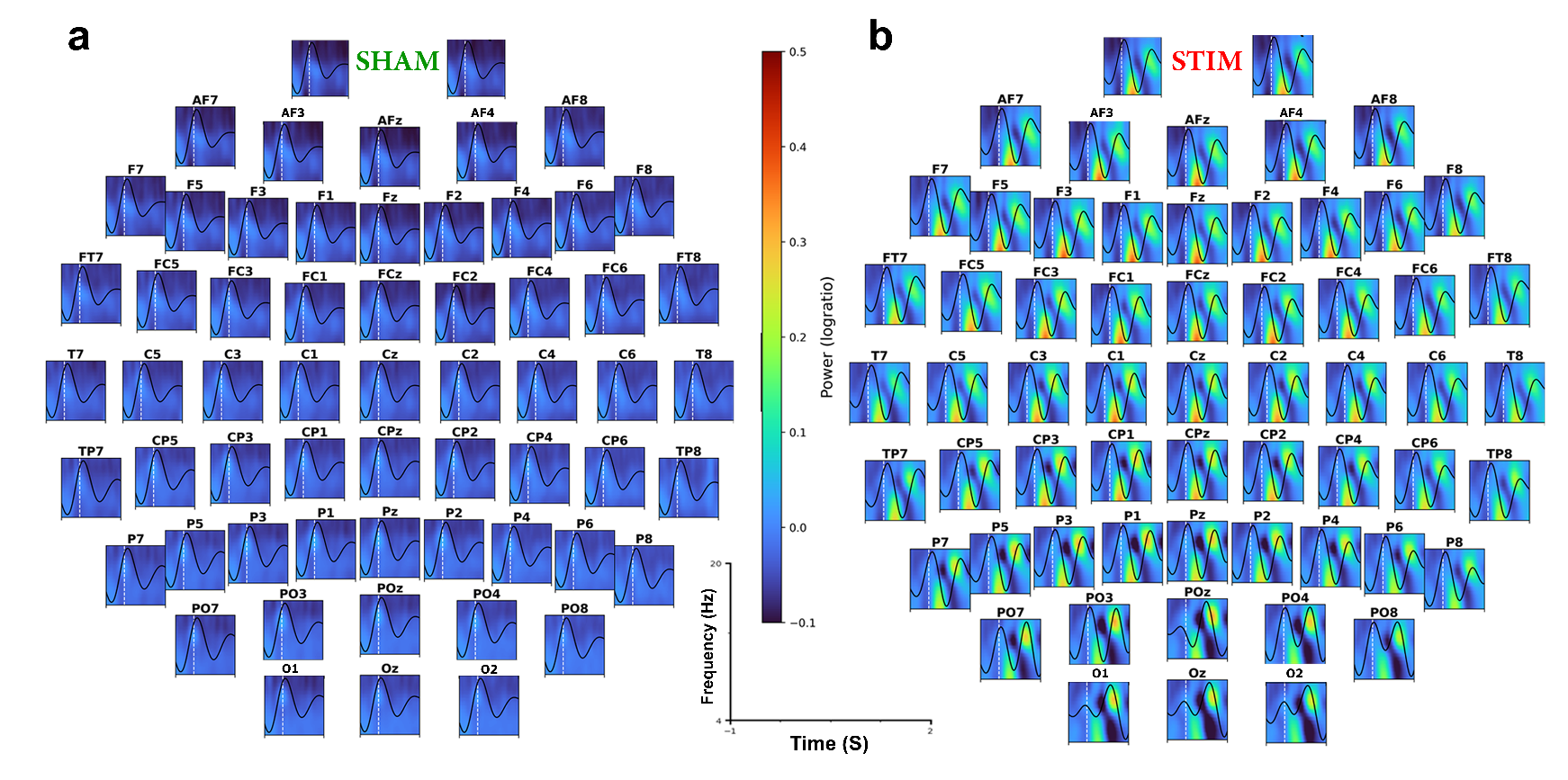

**Figure S4. Each panel displays the TFR for an individual EEG channel arranged according to scalp topography for SHAM (a) and STIM (b) conditions.** Power is expressed as log-ratio relative to the pre-stimulus baseline. The black waveform overlaid on each panel represents the averaged SO ERP, and the dashed vertical line indicates stimulus onset.

**Baseline Sleep Oscillation Characteristics and Responsiveness to Auditory Stimulation**

To examine whether baseline sleep oscillation characteristics predicted responsiveness to auditory stimulation, spectral power during SHAM nights was quantified from NREM sleep EEG recordings. Power spectral density (PSD) estimates were computed using Welch’s method in MNE-Python (compute_psd) with a frequency range of 0.4–30 Hz. For each participant, PSD values (dB µV²/Hz) were averaged across channels and frequencies within each ROI. Baseline oscillatory measures were computed separately for SHAM1 and SHAM2 nights and subsequently averaged to obtain a single baseline estimate per participant. To quantify responsiveness to stimulation, a stimulation benefit score was calculated for each participant as the mean overnight percentage change in word-pair memory performance across the two STIM nights.

Pearson correlation analyses revealed no significant associations between baseline oscillatory activity during SHAM nights and stimulation-related memory benefit (SO: r(14) = −0.213, p = .428; theta: r(14) = −0.302, p = .256; fast sigma: r(14) = −0.186, p = .491), indicating that baseline NREM oscillatory power did not significantly predict responsiveness to auditory stimulation.

Grand-mean topographical maps (Figure S5) of SHAM-night PSD revealed the expected spatial distributions of oscillatory activity across the scalp. Slow oscillation power was maximal over frontal regions, and fast sigma activity showed stronger centroparietal expression.

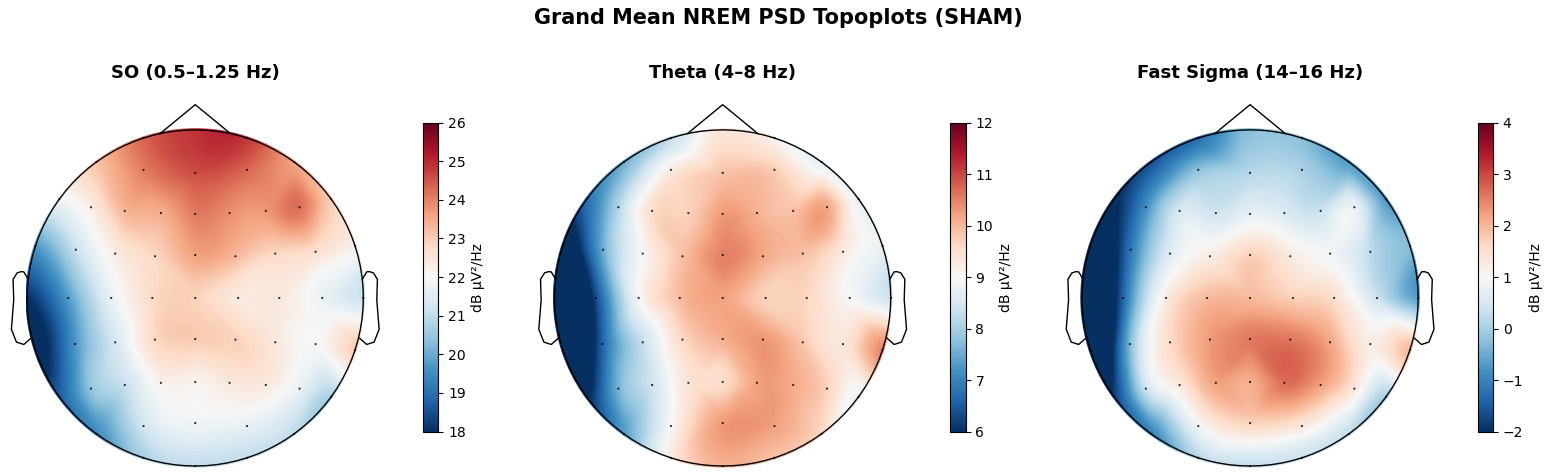

**Figure S5. Grand-mean topographical distributions of SO, theta, and fast sigma power during SHAM nights.**

Overall, these analyses suggest that individual differences in baseline NREM slow oscillation, theta, and fast sigma power did not significantly predict memory responsiveness to closed-loop auditory stimulation in the present sample.

**Do Baseline Sleep Oscillations Predict Neural Responsiveness to PLAS?**

To determine whether baseline sleep oscillatory activity predicted neural responsiveness to stimulation, PLAS response was computed as the difference between STIM and SHAM power values. Pearson correlation analyses were subsequently performed between baseline oscillatory power measured during SHAM nights and the corresponding stimulation-induced change scores.

No significant associations were observed between baseline slow oscillation or fast sigma power and stimulation-induced changes in the corresponding oscillatory activity (SO: r(14) = .071, p = .794; fast sigma: r(14) = .085, p = .755). In contrast, baseline theta power was significantly negatively associated with stimulation-induced theta enhancement, r(14) = −.568, p = .022). This result suggest that participants with lower baseline theta activity showed larger theta increases during stimulation (Figure S6).

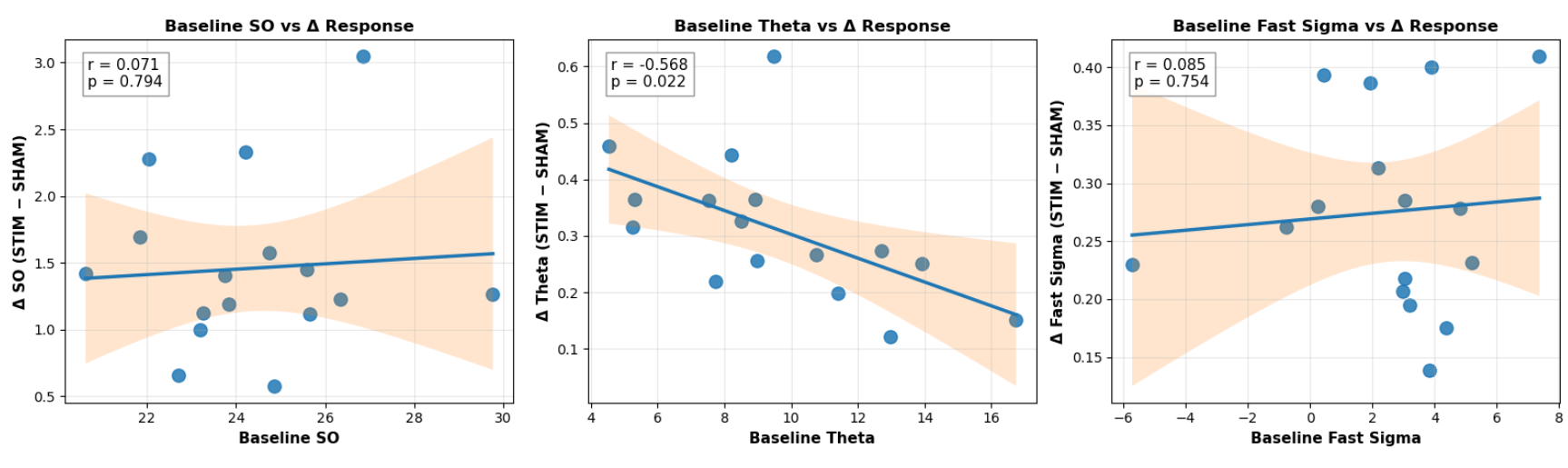

**Figure S6. Relationships between baseline power measured during SHAM nights and stimulation-induced changes in oscillatory activity (STIM − SHAM).**
